# The conserved LEM-3/Ankle1 nuclease is involved in the combinatorial regulation of meiotic recombination repair and chromosome segregation in *Caenorhabditis elegans*

**DOI:** 10.1101/223172

**Authors:** Ye Hong, Maria Velkova, Nicola Silva, Marlène Jagut, Viktor Scheidt, Karim Labib, Verena Jantsch, Anton Gartner

**Affiliations:** Centre for Gene Regulation and Expression, University of Dundee, United Kingdom; Department of Chromosome Biology, Max F. Perutz Laboratories, University of Vienna, Austria; MRC Protein Phosphorylation and Ubiquitylation Unit, School of Life Sciences, University of Dundee, Dundee, UK

## Abstract

Homologous recombination is essential for crossover (CO) formation and accurate chromosome segregation during meiosis. It is of considerable importance to work out how recombination intermediates are processed leading to CO and non-crossover (NCO) outcome. Genetic analysis in budding yeast and *Caenorhabditis elegans* indicates that the processing of meiotic recombination intermediates involves a combination of nucleases and DNA repair enzymes. We previously reported that in *C. elegans* meiotic Holiday junction resolution is mediated by two redundant pathways, conferred by the SLX-1 and MUS-81 nucleases, and by the HIM-6 Blooms helicase in conjunction with the XPF-1 endonucleases, respectively. Both pathways require the scaffold protein SLX-4. However, in the absence of all these enzymes residual processing of meiotic recombination intermediates still occurs and CO formation is reduced but not abolished. Here we show that the LEM-3 nuclease, mutation of which by itself does not have an overt meiotic phenotype, genetically interacts with *slx-1* and *mus-81* mutants, the respective double mutants leading to 100% embryonic lethality. LEM-3 and MUS-81 act redundantly, their combined loss leading to a reduced number of early meiotic recombination intermediates, to a delayed disassembly of foci associated with CO designated sites, and to the formation of univalents linked by SPO-11 dependent chromatin bridges (dissociated bivalents). However, LEM-3 foci do not co-localize with ZHP-3 a marker that congresses into CO designated sites. In addition, neither CO frequency nor distribution is altered in *lem-3* single mutants or in combination with *mus-81* or *slx-4* mutations, indicating that LEM-3 drives NCO outcome. Finally, we found persistent chromatin bridges during meiotic divisions in *lem-3; slx-4* double mutants. Supported by the localization of LEM-3 between dividing meiotic nuclei, this data suggests that LEM-3 is able to process erroneous recombination intermediates that persist into the second meiotic divisions.

**Author Summary:** Meiotic recombination is required for genetic diversity and for the proper chromosome segregation. Recombination intermediates, such as Holliday junctions (HJs), are generated and eventually resolved to produce crossover (CO) and non-crossover (NCO). While an excess of meiotic double-strand breaks is generated, most breaks are repaired without leading to a CO outcome and usually only one break for each chromosome pair matures into a CO-designated site in *Caenorhabditis elegans*. Resolution of meiotic recombination intermediates and CO formation have been reported to be highly regulated by several structure-specific endonucleases and the Bloom helicase. However, little is known about enzymes involved in the NCO recombination intermediate resolution. In this study, we found that a conserved nuclease LEM-3/Ankle1 acts in parallel to the SLX-1/MUS-81 pathway to process meiotic recombination intermediates. Mutation of *lem-3* has no effect on CO frequency and distribution, indicating LEM-3 functions as a nuclease promoting NCO outcome. Interestingly, a prominent localization of LEM-3 is found between dividing meiotic nuclei. We provide evidence that LEM-3 is also involved in processing remaining, erroneous recombination intermediates during meiotic divisions.

## Introduction

Meiosis is comprised of two specialized cell divisions that elicit the reduction of the diploid genome to haploid gametes. Homologous recombination occurs in the first meiotic division and is required for meiotic crossover (CO) formation [1]. COs are needed to shuffle genetic information between maternal and paternal chromosomes and are thus required to ensure genetic diversity. COs become cytologically visible as chiasmata and also provide stable connections between maternal and paternal homologous chromosomes (homologs). Chiasmata counter the spindle force and thereby facilitate the accurate segregation of homologs in the first meiotic division.

Meiotic recombination is initiated by DNA double-strand breaks (DSBs) generated by the conserved meiosis-specific Spo11 protein [2]. The number of DSBs generated by SPO11 exceeds the number of COs, ranging from ~2:1 in *Saccharomyces cerevisiae* to ~20:1 in maize [3–6]. In *Caenorhabditis elegans* typically only one DSB per homologous pair will mature into a CO event [7]. It is thought that the excessive number of DSBs is required to ensure that at least one CO happens on each homolog, a notion supported by checkpoint mechanisms that delay meiotic prophase progression when the number of DSBs are reduced [8–10]. It is unclear how CO is selected from the pool of DSBs. The CO selection (or designation) correlates with the congression of several pro-CO factors into six distinct foci, one on each paired chromosome starting from the mid-pachytene stage. These include the cyclin-related protein COSA-1/CNTD1, MSH-5 a component of the MutSY, the predicted ubiquitin ligases ZHP-3/RNF212 and HEI10, the BLM helicase HIM-6 as well as its regulatory subunit RMH-1 [11–16].

When the CO designation sites are stabilised and protected by those pro-CO factors, processing of meiotic recombination intermediates is biased towards the CO outcome. One of the meiotic recombination intermediates is called a Holliday junction (HJ), a cruciform DNA structure formed as a result of a reciprocal exchange DNA strands between homologous chromosomes [17]. While in fission yeast single HJs appear to predominate, direct evidence for the occurrence of double HJs (dHJs) was obtained in budding yeast [18]. dHJs can be processed to result in CO or a non-crossover (NCO) outcome, depending on the way the cut is made by structure-specific endonucleases [19]. There is emerging evidence that a combination of nucleases is required for the processing of meiotic HJs to promote CO formation [20]. Only in fission yeast, deletion of a single nuclease, MUS81, leads to a defect in meiotic CO formation [21]. In budding yeast, absence of the MUS81-MMS4, SLX1-SLX4 or GEN1 nucleases exhibits a modest reduction of meiotic COs [22]. The Exo1 nuclease and the mismatch-repair MutLγ complex Mlh1-Mlh3 have also been shown to contribute to HJ resolution [22, 23]. Mouse *gen1* mutants have no meiotic phenotype, while *mus81* animals only show minor phenotypes [24]. In *C. elegans* HJ resolution and CO formation appear to be conferred by at least two redundant pathways [25–27]. One pathway is defined by the MUS-81 and SLX-1 nucleases. Consistent with *in vitro* nuclease assays, it appears that SLX-1 might confer a first nick on a HJ, the nicked HJ being the preferred substrate for MUS-81 [28]. The second pathway comprises the XPF-1 nuclease and the Bloom (BLM) helicase HIM-6. It is possible that HIM-6 might be able to unwind a HJ, which would generate a substrate cleaved by XPF-1. Both pathways require SLX-4 as a scaffold protein. When both pathways are compromised, the CO frequency is reduced by roughly one third [25].

Since only a small subset of DSBs are designated as CO sites, the majority of DSBs have to be processed in favour of the NCO outcome [29]. In budding yeast recombination sites leading to NCO outcome mature early and are separated from CO designated sites during early pachytene stage [30, 31]. Several helicases are proposed to suppress COs by disassembly of early recombination intermediates such as D-loop structure, and promote NCO formation in a pathway called synthesis-dependent strand annealing (SDSA). In budding yeast this is driven by the Srs2 helicase [32]. In animals, this activity is ascribed to the BLM and RTEL helicases. In *C. elegans* deletion of the RTEL-1 helicase leads to an elevated number of meiotic COs [33]. In contrast, deletion of *him-6*, the *C. elegans* BLM homologue, leads to a reduced number of meiotic CO formation and a late pro-CO function at CO designation sites was recently implied [34, 35]. Once dHJs are formed, they can either be dissolved by the BLM helicase and Top3 topoisomerase in a NCO manner or resolved by nucleases to form CO or NCO [32].

In this study, we report on roles of the LEM-3/Ankle1 nuclease in processing meiotic recombination intermediates. LEM-3 is only conserved in animals and the mammalian ortholog is referred to as Ankle1 [36, 37]. *C. elegans lem-3* mutants are hypersensitive to ionizing irradiation, UV treatment and DNA cross-linking agents [36]. LEM-3/Ankle1 contains an N-terminal LEM domain, Ankyrin repeats and a GIY-YIG nuclease motif. The same nuclease motif can also be found in bacterial UvrC nucleotide excision repair proteins and in the distantly related SLX1 nuclease [38]. Our data show that LEM-3 and MUS-81 act redundantly to process early recombination intermediates during meiosis. Loss of LEM-3 and MUS-81 leads to aberrant profiles of recombination markers, delayed processing of markers for CO designation, increased apoptotic cells and formation of dissociated bivalents. In addition, we found that a considerable pool of LEM-3 localizes between dividing meiotic nuclei and chromosome segregation is compromised due to persistent chromatin linkages in the absence of both LEM-3 and SLX-4, indicating that LEM-3 is able to process erroneous recombination intermediates that persist into meiotic divisions.

## Results

### LEM-3 acts in a genetic pathway parallel to SLX-4 and MUS-81/SLX-1

We and other groups previously showed that there might be two pathways needed for the resolution of meiotic recombination intermediates, one dependent on SLX-1 and MUS-81 and the other one needing XPF-1, while SLX-4 is acting in both pathways [25–27]. Given that viability and CO recombination is reduced but not eliminated when both the MUS-81 and XPF-1 pathways are compromised, we suggested that there might be at least one additional nuclease that has not been identified. We therefore searched for nucleases which are synthetic lethal with SLX-4 and focused on LEM-3 in this study. Out of the three previously reported LEM-3 alleles, we used the *lem-3 (mn155)* and the *lem-3 (tm3468)* alleles, the former leads to a premature stop codon at amino acid 190 leaving the N-terminal Ankyrin Repeat domain intact, but eliminating the nuclease domain, thus representing an null allele [36]. *lem-3 (tm3468)* causes a in frame 110 amino acid deletion between the Ankyrin Repeat and the LEM domain [36]. We found that the viability of *lem-3 (tm3468); slx-4* was reduced to ~10% while *lem-3 (mn155); slx-4* worms were 100% embryonic lethal. Given that *lem-3 (mn155)* is a null allele we focused our genetic analysis on that allele and generated double mutants with *slx-1, mus-81 and xpf-1*. We found 100% lethality with *slx-1 lem-3*, and *mus-81 lem-3* double mutants (Figure 1). In contrast, the viability of *xpf-1* was not further reduced in *lem-3 xpf-1* double mutants (Figure 1). In summary, these initial genetic data indicate that LEM-3 might act in parallel to SLX-4. LEM-3 could also be involved in the XPF-1 pathway but in parallel to the SLX-1/MUS-81 pathway.

**Figure 1.**
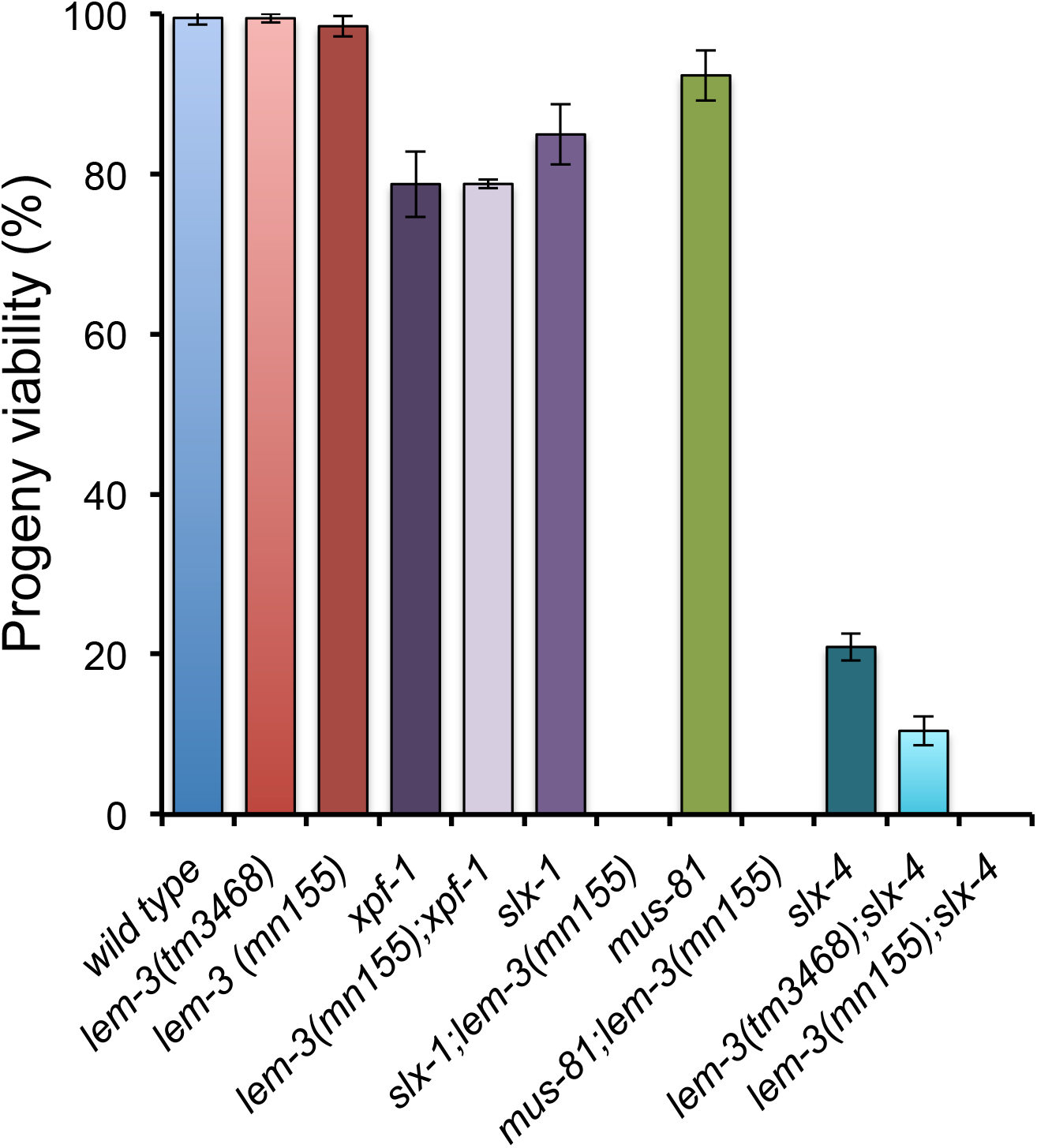
Genetic interaction between LEM-3, MUS-81, SLX-1 and SLX-4 endonucleases. Progeny viability in % was determined by counting number of viable eggs/total number of eggs laid. Error bars represent standard deviation of the mean.

### Meiotic chromosome axis formation and CO designation are normal in *lem-3* mutants

We next wanted to test if the synthetic phenotypes we observed are linked to defects in meiotic prophase progression. In *C. elegans* meiotic prophase progression, which occurs in a gradient of differentiation, can be visualized using dissected germ lines. At the distal end of the gonad germ cell mitotically divide, before entering the transition zone where meiotic chromosomes reorganise into arrays of chromatin loops anchored to the chromosome axis [39]. The chromosome axis establishes a platform for homologous chromosome paring, DSB induction, synaptonemal complex assembly, and CO formation [40]. Staining with HTP-3 a component of axial element indicated that chromosome axis formation occurred normally in *lem-3 and slx-4 single mutants and lem-3; slx-4* double mutants during pachytene stage (Figure S1A).

We also tested whether CO designation occurs normally using a strain expressing a functional COSA-1::GFP fusion in *mus-81 lem-3* and *lem-3; slx-4* double mutant worms. COSA-1 foci mark CO designated sites during late pachytene [11]. As previously reported for wild type, *slx-4* and *mus-81* single mutants, only 6 CO designated sites are apparent in the *lem-3* single mutant and *mus-81 lem-3* and *lem-3; slx-4* double mutants in late pachytene (Figure S1B and S1C), indicating that CO designation is not perturbed.

### LEM-3 is required for meiotic recombination intermediate processing

Since there are no overt defects in chromosome axis formation and CO designation, we assessed if the synthetic lethal phenotypes we observed were due to a defect in meiotic recombination. We monitored pyknotic cells that have abnormally condensed nuclei and found that they became apparent in mid/late pachytene in the *mus-81* single mutant and to a larger extent in the *mus-81 lem-3* double mutant (Figure 2A). Directly scoring for the number of apoptotic corpses by DIC (differentia interference contrast) microscopy revealed that apoptosis was increased in *mus-81* worms, further increased in *slx-4* worms, and that the highest level of apoptosis occurred in both *mus-81 lem-3* and *lem-3; slx-4* double mutant worms (Figure 2B).

**Figure 2.**
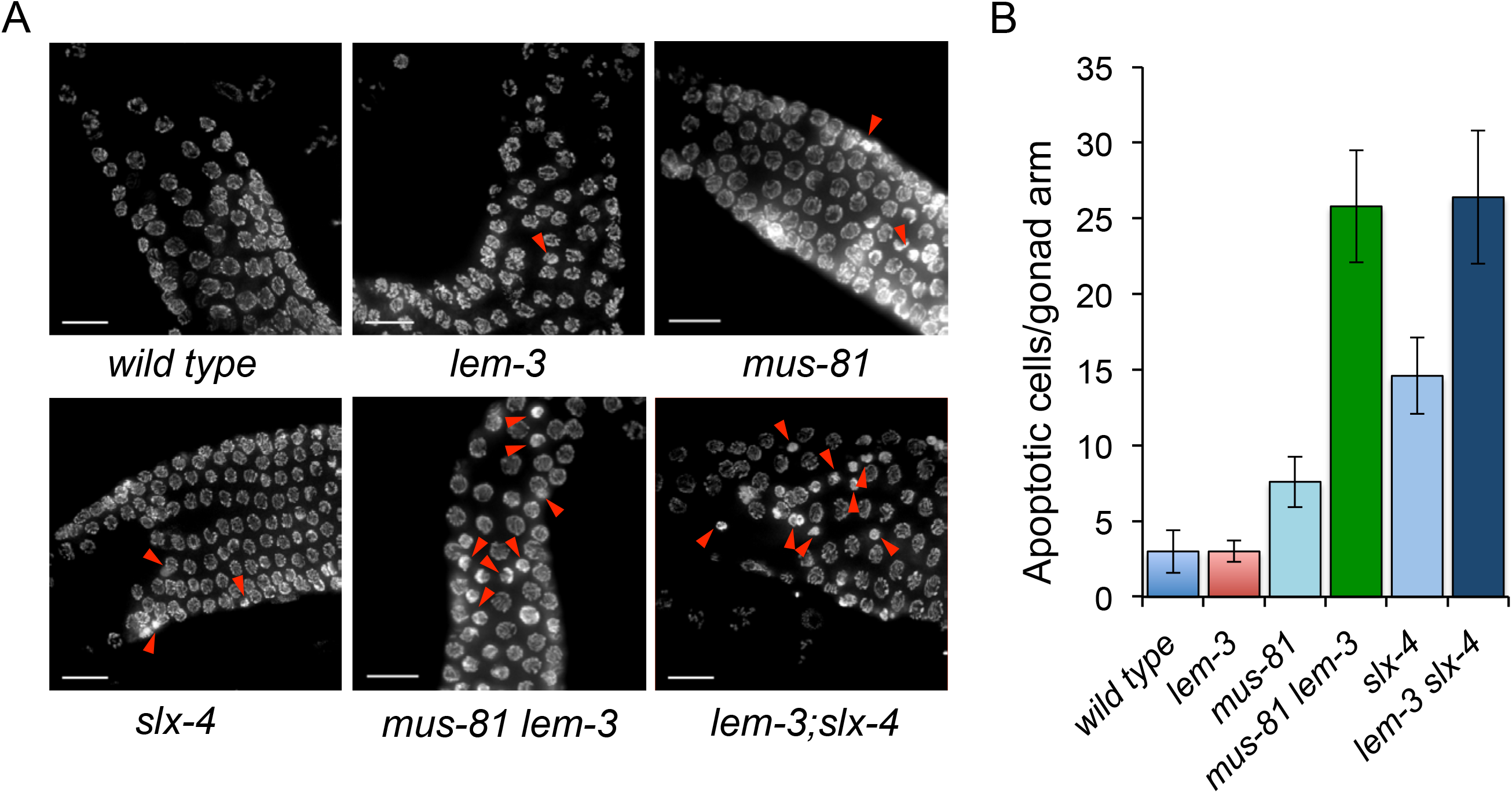
Mutation of *lem-3* in *mus-81* and *slx-4* mutants cause increased apoptosis. (A) Representative images of DAPI-stained germline in wild type, *lem-3, mus-81, slx-4; mus-81 lem-3; lem-3; slx-4* mutants. Red arrowheads indicate pyknotic cells with abnormally condensed nuclei in pachytene stage. Scale bars: 15 μm. (B) Quantification of apoptotic cells per gonad in the indicated genotypes. Error bars represent standard deviation of the mean.

While an excess of meiotic DSBs are generated, most breaks are repaired without leading to a CO outcome and only one break for each chromosome pair matures into a CO-designated site in *C. elegans* [41]. RMH-1 is a marker, which labels both CO and NCO recombination intermediates [16]. Consistent with the above-mentioned epistasis experiments both *lem-3* and *mus-81* single mutants showed a wild type distribution of RMH-1 foci in early pachytene (with an average of 11 foci) and predominantly 6 foci in late pachytene. Slightly higher numbers of foci were seen in mid-pachytene in *lem-3* (on average14.8 foci) and *mus-81* (12.9), compared to 10.8 in the wild type (Figure 3). In contrast, in the *mus-81 lem-3* double mutant foci numbers dropped to an average of 7.5 foci in early pachytene, 8.2 in mid-pachytene and 4.6 foci in late pachytene (Figure 3). A similar trend can be observed for RAD-51 profiles. RAD-51 foci mark on-going recombination intermediates. The number of RAD-51 foci in pachytene is close to wild type in *mus-81 lem-3* double mutants worms, while RAD-51 foci are increased in *mus-81* and *lem-3* single mutants (Figure 6B). In summary our results are consistent with the number of early recombination intermediates being reduced in *mus-81 lem-3* double mutants as compared to *lem-3* and *mus-81* single mutants.

**Figure 3.**
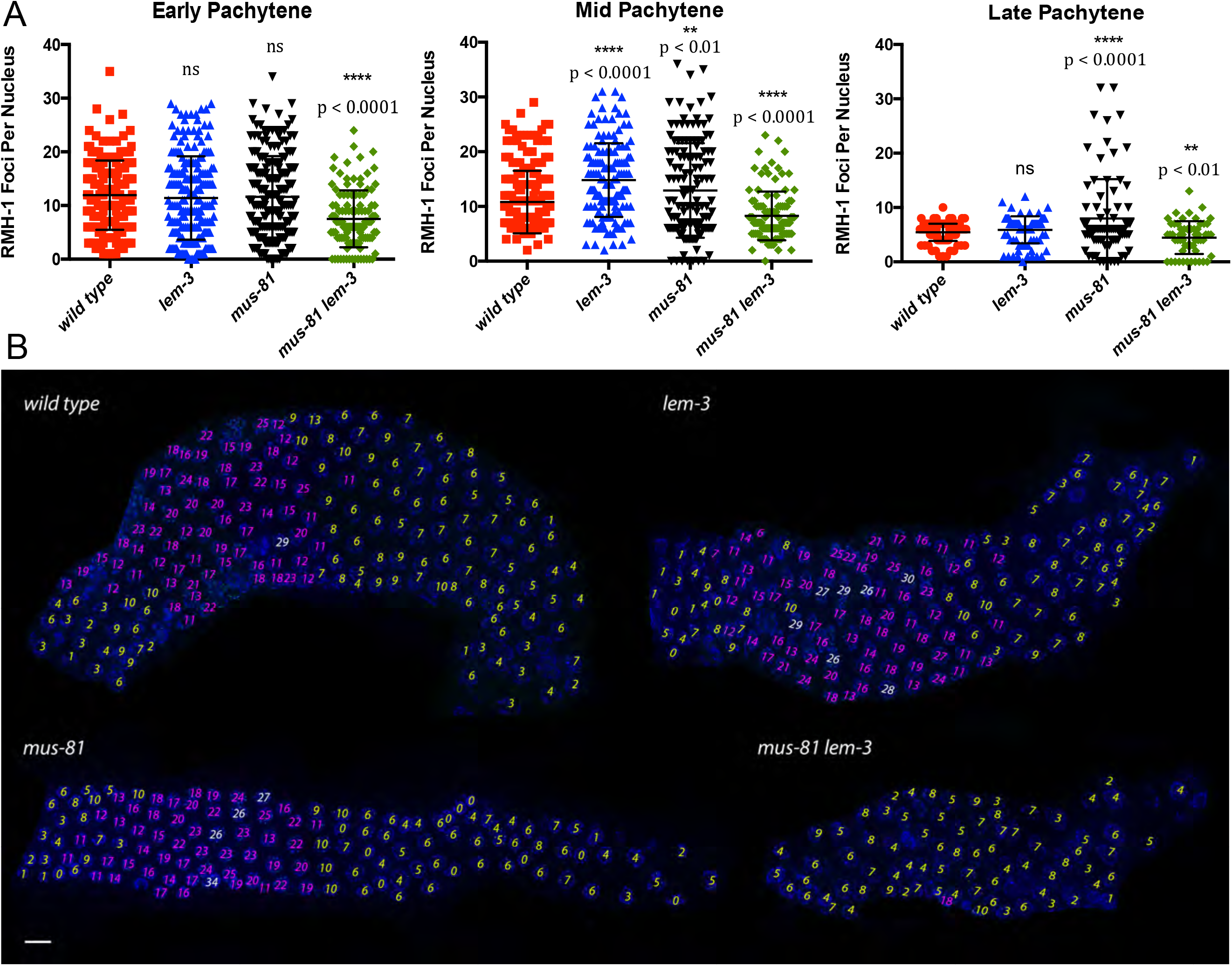
Comparison of RMH-1::GFP localization in wild type, *lem-3, mus-81* single mutants and *mus-81 lem-3* double mutants. (A) Quantification of RMH-1::GFP foci counts in wild type, *lem-3, mus-81* and *mus-81 lem-3* mutants in early Pachytene, mid Pachytene and late Pachytene stages. Quantifications were done for three gonads per genotype. Asterisks indicate statistical significance as determined by student T test. P Values below 0.05 were considered as significant, p < 0.05 is indicated with *, p < 0.01 with ** and p < 0.0001 with ****. (B) Representative images of gonads stained with DAPI. Yellow numbers represent those nuclei with RMH-1::GFP foci between 1 and 10, pink numbers represent nuclei with RMH-1::GFP counts between 11 and 25, and white numbers represent nuclei with RMH-1::GFP foci above 25.

**Figure 4.**
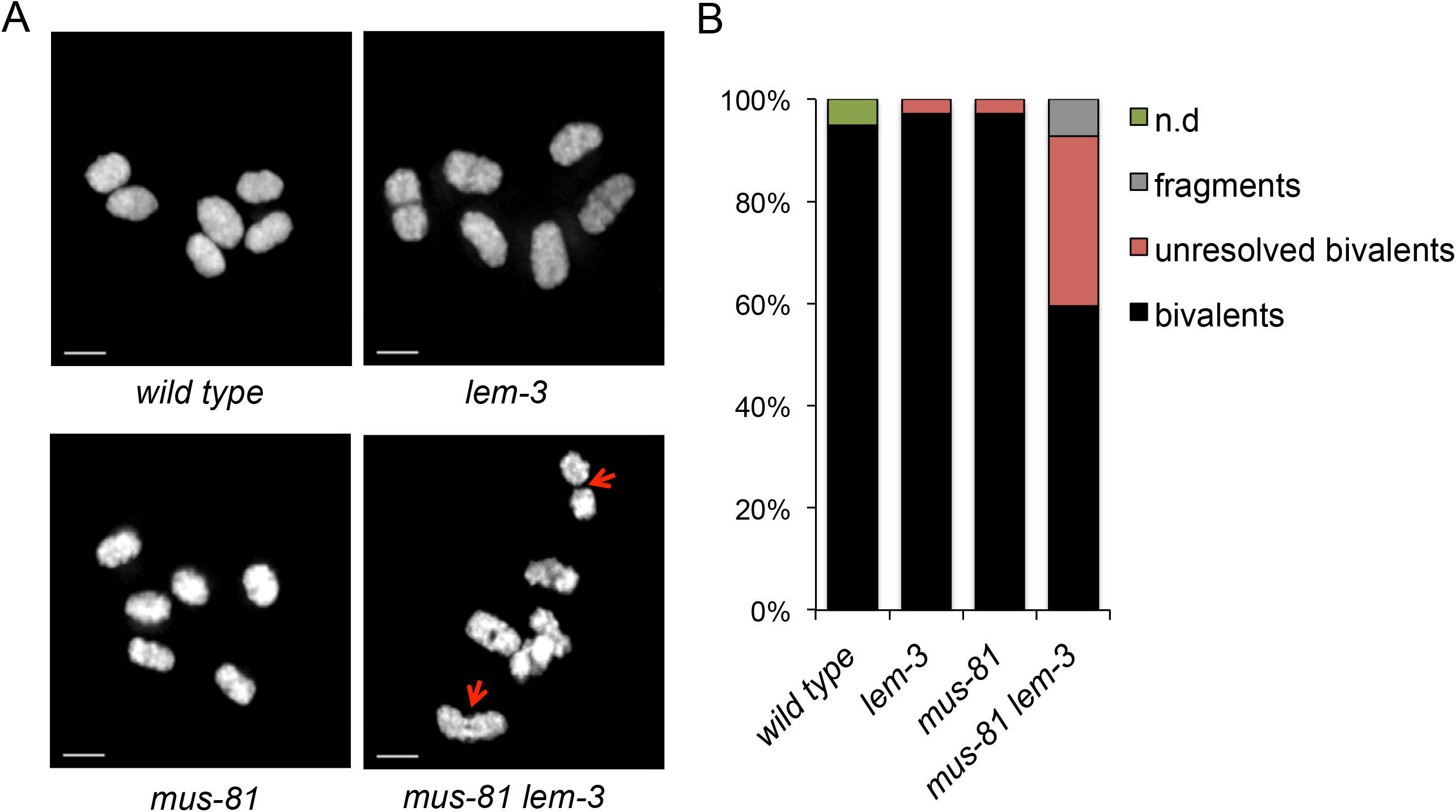
Depletion of LEM-3 and MUS-81 leads to formation of dissociated bivalents. (A) Images of DAPI-stained chromosomes in −1 oocytes at diakinesis in wild type, *lem-3, mus-81* and *mus-81 lem-3* mutants. Red arrows indicate dissociated bivalents. Scale bars: 2 μm. (B) Quantification of bivalents, ‘dissociated bivalents’ and fragments observed in indicated genotypes. Overlapping chromosomes that could not be assigned to the above categories were scored as “n/d”.

**Figure 5.**
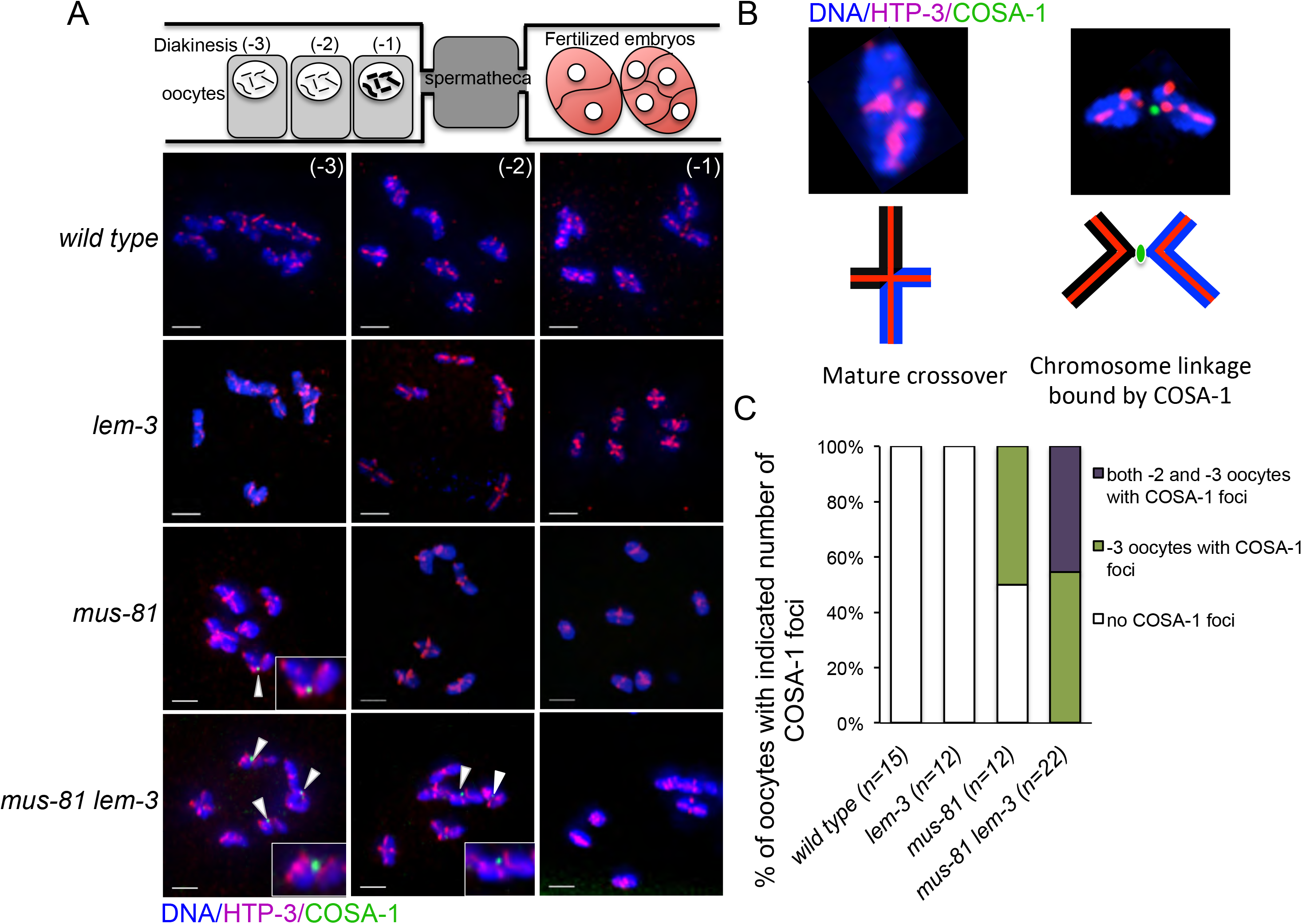
Delayed crossover maturation in *mus-81 lem-3* double mutants. (A) DAPI staining of representative diakinesis nuclei of wild type, *lem-3, mus-81* and *mus-81 lem-3* mutants. White arrowheads indicate the remaining GFP::COSA-1 foci. (B) Scheme depicting unresolved crossover recombination intermediate bound by COSA-1 at the diakinesis stage (C) Quantification of oocytes with indicated number of COSA-1 foci. The sample size (n) indicates the number of germline examined for each genotype.

**Figure 6.**
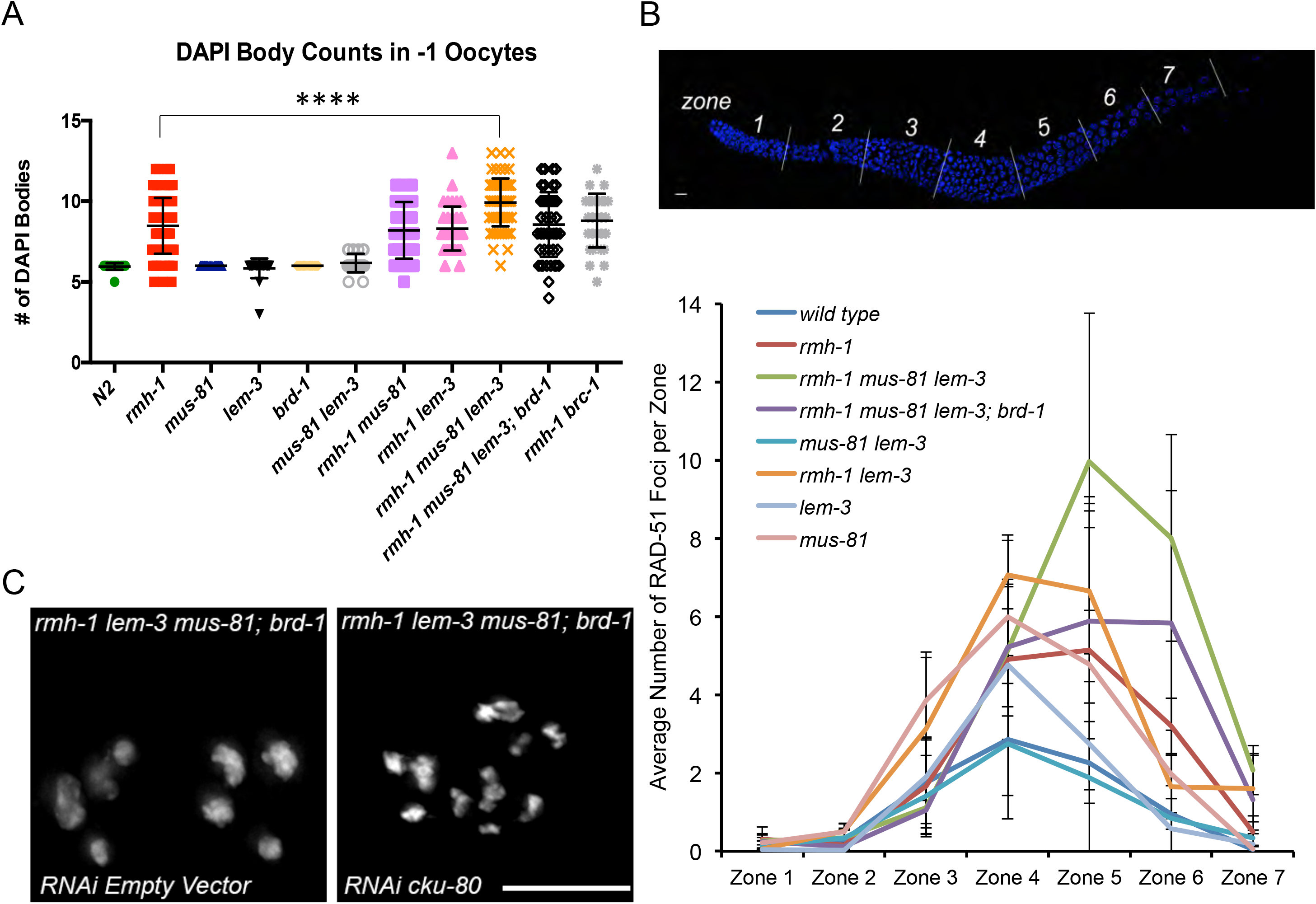
Univalents in *rmh-1 mus-81 lem-3* triple mutants can be rescued by depletion of *brd-1*. (A) Quantification of DAPI-stained bodies in −1 oocytes at diakinesis in the indicated genotypes. The sample size (n) are indicated as follows: wild type (n=23), *rmh-1* (n=88), *mus-81* (n=32), *lem-3* (n=26), *brd-1* (n=30), *mus-81 lem-3* (n=23), *rmh-1 mus-81* (n=46), *rmh-1 lem-3* (n=40), *rmh-1 mus-81 lem-3* (n=73), *rmh-1 mus-81 lem-3; brd-1* (n=50), *rmh-1 mus-81; xpf-1* (n=35), *rmh-1 lem-3; xpf-1* (n=38). (B) Quantification of RAD-51 profiles over the course of meiotic prophase. *C. elegans* gonads were divided in seven equal zones. These were then counted and quantified separately for RAD-51. Quantifications were done based on three representative gonads per genotype. (C) Representative images of DAPI stained −1 oocytes in *rmh-1 lem-3 mus-81; brd-1* mutant treated with empty vector and *cku-80* RNAi.

The morphology and number of diakinesis chromosomes can serve as readouts of meiotic recombination defects. During diakinesis, homologous chromosome pairs restructure to form bivalents, which can be observed as 6 DAPI-stained bodies in wild-type maturing oocytes [7]. Defects in meiotic recombination can result in a failure to stably connect homologous chromosomes, which becomes apparent as univalents at diakinesis (12 when physical linkages between all six homologue pairs fail to form). Our prior analysis of *slx-4*, as well as *mus-81; xpf-1* double mutants revealed a distinct phenotype [25]. In contrast to *spo-11*, pairs of ‘univalents’ were found associated with each other, being linked by SPO-11 dependent chromatin bridges. We termed these structures as ‘dissociated bivalents’. We interpreted these structures as chromosome pairs that engage in meiotic recombination but do not resolve recombination intermediates thus leading to the linkage of maternal and paternal chromosomes. As expected, wild type, *lem-3*, and *mus-81* single mutants predominately contained 6 bivalents (Figure 4). In contrast, *mus-81 lem-3* double mutant showed increased number of dissociated bivalents, which can also be detected in *slx-1 lem-3* mutant (Figure 4 and S2). Importantly, the analysis of *slx-1 lem-3; spo-11 triple* mutants revealed 12 univalents (Figure S2), indicating that the chromosome linkages we observed are *spo-11* dependent and thus represent meiotic recombination intermediates.

In the wild type, COSA-1 foci represent CO precursors and develop as strong foci at late pachytene before gradually dissociating from chromosome pairs through diplotene and diakinesis [11]. While COSA-1 foci are fully gone in wild type and *lem-3* single mutant prior to the −3 oocyte stage (Figure 5A), they could still be detected in 50% of −3 oocytes of the *mus-81* single mutant *(n=12)*. COSA-1 foci persisted up to 45% of the −2 oocytes in *mus-81 lem-3* double mutant (n=22), but finally disappeared in −1 oocytes (Figure 5C). The COSA-1 foci localized on the chromosome linkages between dissociated bivalents could be eventually resolved to form CO or NCOs (Figure 5B). Taken together, these data suggest that compromised recombination intermediate processing in *mus-81 lem-3* double mutant leads to a delay in dismantling COSA-1 foci.

### Evidence for LEM-3 working as an NCO resolvase

The rate of CO recombination is reduced in *slx-4* single and *mus-81; xpf-1* double mutants by approximately one third. Given that *slx-1 lem-3, mus-81 lem-3* and *lem-3; slx-4* double mutants only sire dead embryos, we expected that CO recombination might be abolished in those double mutants. CO frequency and distribution can be investigated by meiotic recombination mapping [42]. We generated the *lem-3* single and double mutants with chromosome V being heterozygous for the Hawaii and N2 backgrounds. To score for recombination frequency and distribution we employed five single nucleotide polymorphisms (snip-SNPs), which together cover 92% of chromosome V [25]. Given the lethality of various *lem-3* double mutants, embryos were used for recombination mapping to avoid biasing the analysis on the basis of viability. We found that CO recombination rates are comparable to wild type in *lem-3* single mutants (Figure S3). Furthermore, when analysing the respective compound mutants, we found that *lem-3* did not lead to a decreased CO rate in conjunction with *mus-81* or *slx-4* single mutants (Figure S3). Thus, despite the chromatin linkages that occur in various *lem-3* double mutants, CO recombination is not reduced. We therefore postulate that LEM-3 is not required for interhomolog CO formation.

We next investigated if LEM-3 is able to promote NCO outcome and helps to prevent unscheduled recombination. Our analysis of diakinesis structures of a triple mutant of *lem-3, mus-81* and *rmh-1* (RMH-1 works in conjunction with the bloom helicase HIM-6) indicated an increased number of “peculiar” univalent (Figure S4), which were previously also observed in *rmh-1* and *him-6* single mutants [16, 25, 35]. These univalents are special in displaying a ‘bivalent like’ chromosome domain reorganization in ‘CO-distal’ and ‘CO-proximal’ domains that depends on CO initiation via *msh-4/msh-5* and *zhp-3* but does not require CO completion [16, 25, 35]. *rmh-1* single and *rmh-1 mus-81* double mutants showed an average of 8.5 DAPI stained bodies, with clearly discernible univalents (indicated by red arrowheads, Figure S5) compared to an average of 10 in the *rmh-1 mus-81 lem-3* triple mutant (Figure 6A and S5). In contrast both *lem-3 mus-81* and *lem-3 rmh-1* double mutants showed an average of 6 and 8.5 DAPI stained bodies, comparable to wild type and *rmh-1* single mutants, respectively (Figure 6A and S5). Thus, the occurrence of those univalents only seen in the triple mutant might be the result of several parallel recombination pathways being compromised. We next set out to test if these univalents result from the mis-direction of recombination intermediates towards unscheduled inter-sister repair. In *C. elegans*, the BRCA-1 homolog of the mammalian BRCA1 recombination protein has been proposed to be important for inter-sister repair in the germline [43]. Indeed, the *rmh-1 mus-81 lem-3; brd-1* quadruple mutant displays a reduction in univalent formation, such that an average of 8.5 DAPI stained bodies can be detected (Figure 6A and S5). Blocking inter-sister repair also leads to a diminished accumulation of RAD-51, indicative of a reduced HR repair and possibly suggesting the activity of an alternative DSB repair pathway (Figure 6B). We thus tested if the non-homolog end-joining DSB repair pathway has a role, and consistent with this, depletion of KU-80 end joining factor by RNAi led to massive DNA fragmentation in the *rmh-1 mus-81 lem-3; brd-1* mutant (Figure 6C). Altogether, these results suggest that the combined absence of RMH-1 and LEM-3 in *mus-81* might direct recombinational repair towards an unscheduled NCO outcome.

### LEM-3 promotes proper chromosome segregation during meiotic cell division

To investigate if LEM-3 has a role in NCO recombination repair, we first checked LEM-3 localization by using strain expressing a GFP::LEM-3 fusion to stain for LEM-3. Consistent with previous reports [36], we found that LEM-3 localized as dots outside of the nucleus in the mitotic germ cells of wild type worms (Figure 7A). LEM-3 foci (typically no more than one per cell) were occasionally observed in the pachytene area, both inside and out of the nucleus (Figure 7B). These LEM-3 foci didn’t co-localize with the ZHP-3 marker that congresses into CO designated sites (Figure 7C) [44]. Interestingly, careful examination of cells undergoing meiotic divisions revealed that LEM-3 localized between dividing nuclei during meiosis II (Figure 7D). To analyse whether LEM-3 has a role in meiotic chromosome segregation, we performed live cell imaging of the first and second meiotic cell divisions by using an integrated Histone mCherry::H2B fusion. We reasoned that anaphase of meiosis I and II might be affected if a chromosome linkage remains present between two homologues, or sister chromatids, respectively. Chromosome segregation in the *lem-3* single mutant was similar to wild type (Figure 7E, Movies S1 and S2). In contrast, chromosome linkages appeared during the first meiotic division in *slx-4* mutants (Figure 7E), as we had previously reported for double mutants affecting both the SLX-1/MUS-81 and the HIM-6/XPF pathway [25]. However, no visible chromosome linkage could be detected during the second meiotic division in the *slx-4* mutant, consistent with an important role for SLX-4 in resolving inter-homolog recombination intermediates (Figure 7E, Movie S3) [45]. In contrast, *lem-3; slx-4* double mutants also showed extensive chromosome linkage formation in the second meiotic division (Figure 7E, Movie S4), in addition to the meiosis I chromosome segregation defects. These data indicate that LEM-3 might have a role in processing inter-sister recombination intermediates that persist into the second meiotic division.

**Figure 7.**
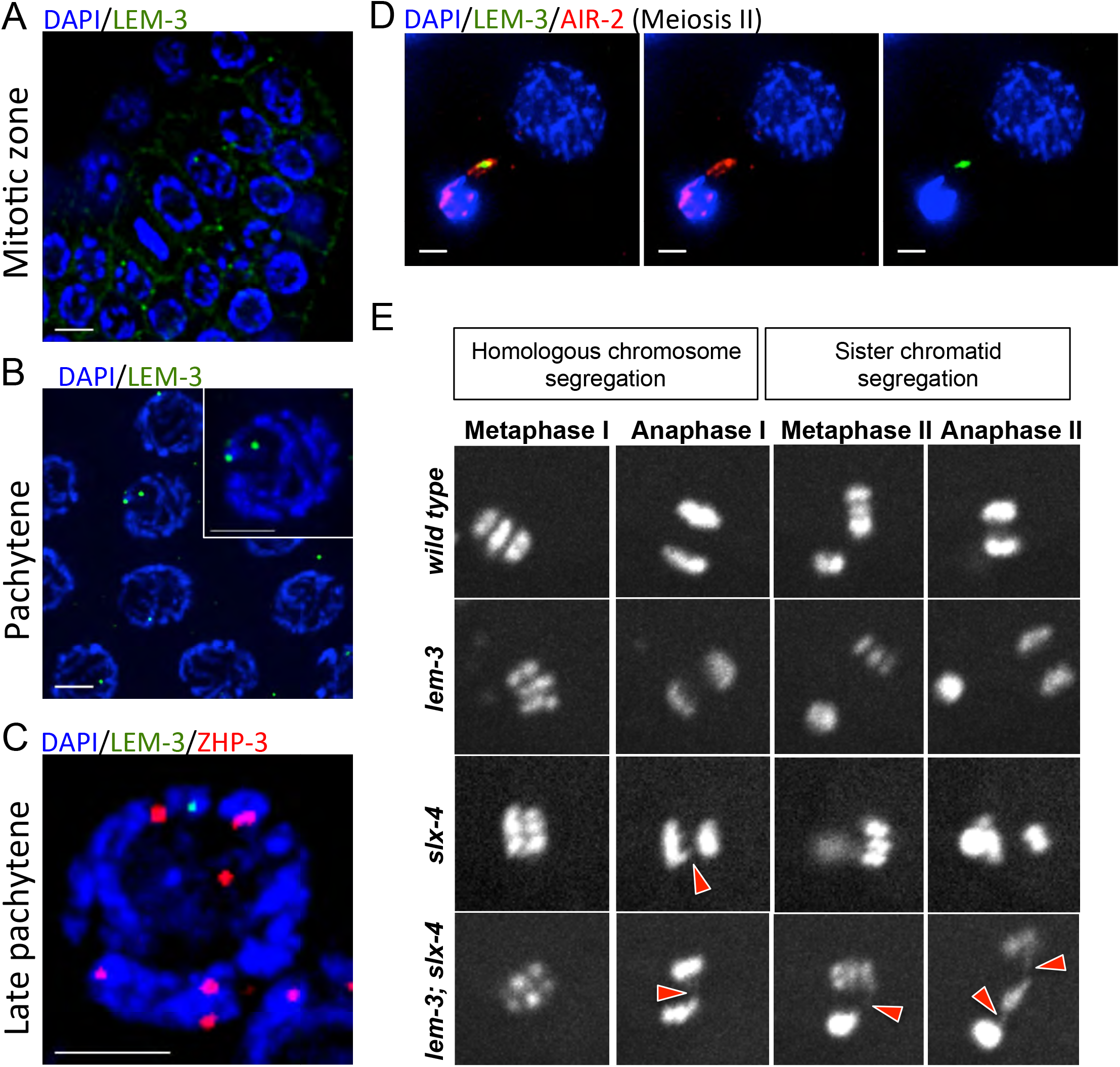
Localization of LEM-3 and its role in meiotic chromosome segregation. Localization of GFP::LEM-3 in mitotic zone (A) and pachytene stage (B). (C) LEM-3 foci (green) do not co-localize with crossover precursor marker ZHP-3 (red). (D) Co-localization of LEM-3 with AIR-2 between dividing nuclei during meiosis II. (E) Representative images taken from time-lapse recordings of mCherry::Histone H2B expressing embryos during meiotic division. Red arrowheads indicate the chromatin linkages.

## Discussion

Timely and efficient resolution of meiotic recombination intermediates requires the combinational action of the BLM helicase and structure-specific endonucleases. Previous studies showed that the MUS81-EME1, SLX1-SLX4, GEN1 and XPF1 nucleases contribute to CO formation in eukaryotes [25–27]. However, removing these nucleases individually or in combination only leads to a reduced but not abolished CO formation and a modest defect in recombination intermediate resolution, indicating the existence of additional enzymes involved in meiotic recombination. In this study, we investigated the interplay between the *C. elegans* LEM-3/ANKLE1 nuclease and the HIM-6 BLM helicase and nucleases previously implicated in meiotic CO resolution. We provide evidence for two roles of LEM-3 during meiosis. Firstly, LEM-3 acts redundantly with the MUS-81 and SLX-1-SLX-4 nucleases to process recombination intermediates. Secondly, LEM-3 functions as a backup nuclease to deal with persistent DNA linkages during meiotic divisions.

The synthetic lethal interaction between LEM-3 and SLX-4 in *C. elegans* led us to investigate whether LEM-3 might act in parallel to the two identified redundant pathways for HJ resolution. Indeed, lacking both LEM-3 and MUS-81 causes an increased number of dissociated bivalents (Figure 4), which represent unresolved recombination intermediates [25]. In addition, CO maturation is delayed in *mus-81 lem-3* double mutants, as revealed by persistent COSA-1 foci at the CO designation sites in −2 oocytes (Figure 5). However, mutation of *lem-3* in combination with *slx-4* or *mus-81* mutants did not lead to a further reduction of CO frequency, or to an altered CO distribution (Figure S3), indicating that LEM-3 might function as an NCO resolvase or a nuclease involved in processing recombination intermediates not destined to become a CO.

RMH-1 foci label both CO and NCO recombination intermediates. A previous study showed that RMH-1 does not colocalize with RAD-51 and appears and disappears later than RAD-51, suggesting that RMH-1 might act after RAD-51 removal and mark late recombination intermediates [16]. We found that the number of both RMH-1 and RAD-51 foci was increased in *lem-3* and *mus-81* single mutants compared to wild type (Figure 3 and 6). The increased steady state number of RMH-1 and RAD-51 recombination intermediates could be due to an increased number of processed DSBs, or due to a delay in DSB processing. However, we found that the *mus-81 lem-3* double mutant has a decreased number of RMH-1 foci in early and mid pachytene compared to wild type (Figure 3). In contrast, the number of RAD-51 foci did not change in the absence of both MUS-81 and LEM-3 (Figure 6), suggesting that MUS-81 and LEM-3 appear to act in conjunction after strand invasion but before the formation of RMH-1 foci. In addition, the number of COSA-1 foci that mark CO designated sites is normal in the *mus-81 lem-3* double mutant (Figure S1). Taken together, our data suggest that LEM-3 and MUS-81 can redundantly process early recombination intermediates and contribute to the formation of recombination intermediates that recruit RMH-1. How might the activity of LEM-3 be involved in processing meiotic recombination intermediates? It has been reported that LEM-3 and its human homolog ANKLE-1 are able to cleave supercoiled plasmid into relaxed circular (nicked) and linear DNA [36, 37]. In addition, LEM-3 can cleave a DNA substrate that is rich in secondary structures [36], indicating that LEM-3 might be a structure-specific endonuclease. Therefore, it is possible that LEM-3 can either process early recombination intermediates, such as D-loops and nicked HJs, or else act at a late stage of meiotic recombination for HJ resolution to generate NCO products. It will be interesting to investigate the DNA substrate preference of LEM-3 in the future.

Chromosome segregation can be affected if unresolved recombination intermediates remain present during meiotic divisions. We previously observed that the extensive chromatin bridges generated by depleting both XPF-1 and MUS-81 pathways during the first meiotic division are eventually resolved in meiosis II, suggesting that backup activities function at or after anaphase I [25]. Here we found that LEM-3 localises between dividing nuclei during meiotic division (Figure 7D). In addition, depletion of LEM-3 leads to accumulation of chromosome linkages in the *slx-4* mutant, especially during meiosis II (Figure 7E), indicating that LEM-3 might have a role in proper chromosome segregation, by directly processing DNA linkages caused by unresolved recombination intermediates. A similar function has also been reported for the Holliday junction resolvase GEN1/YEN1 nuclease in S. *cerevisiae* [46]. The budding yeast *yen1* mutant does not have obvious meiotic defects, whereas the *mus81 yen1* double mutant fails to segregate its chromosomes due to unresolved DNA joint-molecules during meiotic anaphase I and II [47]. Furthermore, the enzymatic activity of YEN-1 is kept at a very low level in prophase but is highly induced at the onset of meiosis II, suggesting that it provides a safeguard activity that processes DNA linkages that escape the attention of MUS81 during meiotic divisions [47]. Mutation of the YEN1/GEN1 nuclease shows phenotypic variation in different organisms [20]. In *C. elegans*, no meiotic phenotypes were observed in the *gen-1* single mutant on its own or in combination with *him-6* or various nuclease mutants [48]. Our data suggest that LEM-3 might provide a failsafe system in *C. elegans* instead of GEN-1, to ensure that all recombination intermediates are resolved at the final stage of gamete formation.

In summary, we provide evidence for a role of LEM-3 in NCO recombination and in mending persistent meiotic recombination intermediates in the second meiotic division. It will be interesting to see whether the mammalian LEM-3 orthologue Ankle1 has a role in meiosis.

## Materials and Methods

### *C. elegans* strains and maintenance

Strains were grown at 20°C followed standard protocols [49]. N2 Bristol was used as the wild type. CB4856 Hawaii was used to generate strains for CO recombination frequency analysis. Strains used in this study are listed in Table S1. The *cop859 [Plem-3::eGFP::STag::lem-3::3’UTRlem-3]* eGFP insertion was generated by Knudra (http://www.knudra.com/) following the procedures described by Dickinson et al [50]. Exact details are available upon request.

### Cytological procedures

Germline immunostaining was performed as described previously with slight modifications [34]. Primary and secondary antibodies were used at the indicated dilutions: rabbit anti-HTP-3 (1:500); guinea pig anti-ZHP-3 (1:250); rabbit anti-AIR-2 (1:200); rabbit anti-RAD-51 (1:1000); mouse anti-GFP (1:500); anti-rabbit Alexa488 (1:400) (Invitrogen), and anti-mouse Alexa488 (1:500) (Invitrogen) and anti-rabbit Alexa Fluor 568 (1:750) (Life technologies). For DAPI staining the final concentration used was 2 μg/μL.

### Recordings of meiotic divisions

Meiotic divisions were recorded by in utero embryo live imaging [51]. Worms were picked into a solution containing 1 mM levamisole to paralyze worms. Worms were mounted on 2% agar pads covered with a coverslip. Images were captured every 10 seconds using spinning-disk confocal microscopy.

### Image acquisition and analysis

Microscopy images acquired with a Delta Vision microscopy were deconvolved and analysed using softWoRx Suite and softWoRx Explorer software (AppliedPrecision, Issaquah, WA, USA). Images acquired with a spinning-disk confocal microscopy were analyzed by ImageJ software.

### Determining meiotic crossover frequency and distribution

Meiotic CO frequency and distribution were assayed essentially as described [25] with slight modifications. Five snip-SNPs on ChrV that differ between N2 Bristol and CB4856 Hawaii were used to determine the crossover landscape in embryos. Single embryos were transferred into lysis buffer by mouth pipetting using a capillary and incubated at −80°C for at least 5 min to help crack the embryo before lysis.

## Acknowledgments

This work was funded by Wellcome Trust Senior Research fellowship (AG 0909444/Z/09/Z), Investigator (KL 102943/Z/13/Z) and Strategic awards (097045/B/11/Z), a MRC core grant KL (MC_UU_12016/13) and the FWF (VJ P28464-B28 and SFB-F34). VS and YH were supported by a Wellcome PhD fellowship and ISSF funds. The Dundee Imaging Facility was supported by the Wellcome Trust (097945/B/11/Z) and the MRC (MR/K015869/1). Thanks to Federico Pelisch, Monique Zetka, Fadri Martinez-Perez, Anne Villeneuve, Abby Dernburg and Kentaro Nabeshima for sharing antibodies. We are grateful to Paul Appleton and Graeme Ball for technical assistance with microscopy, to Shohei Mitani and the Japanese National Bioresource Project for the Nematode for *xpf-1, slx-1* and *mus-81* mutants and the *Caenorhabditis* Genetics Center for providing further strains.

---

**Figure S1.**
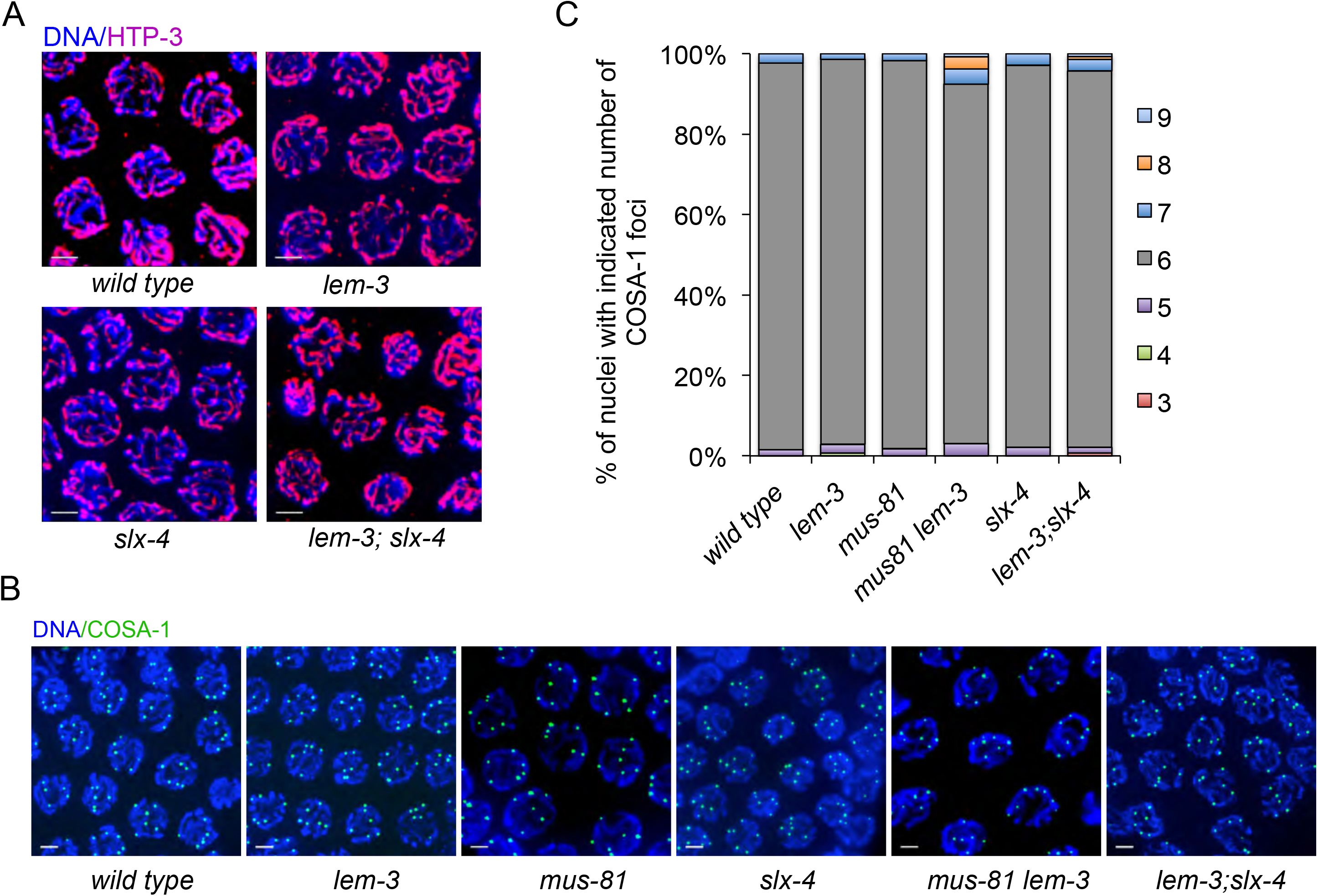
Meiotic chromosome axis formation and crossover designation are normal in *mus-81 lem-3* and *lem-3; slx-4* double mutants. (A) Representative images of pachytene nuclei stained with an antibody recognizing the chromosome axis component HTP-3 (red) and DAPI (blue). Scale bars: 2 μm. (B) DAPI staining of representative pachytene nuclei containing GFP::COSA-1 foci. Scale bars: 2 μm. (C) Quantification of nuclei with indicated number of COSA-1 foci.

**Figure S2.**
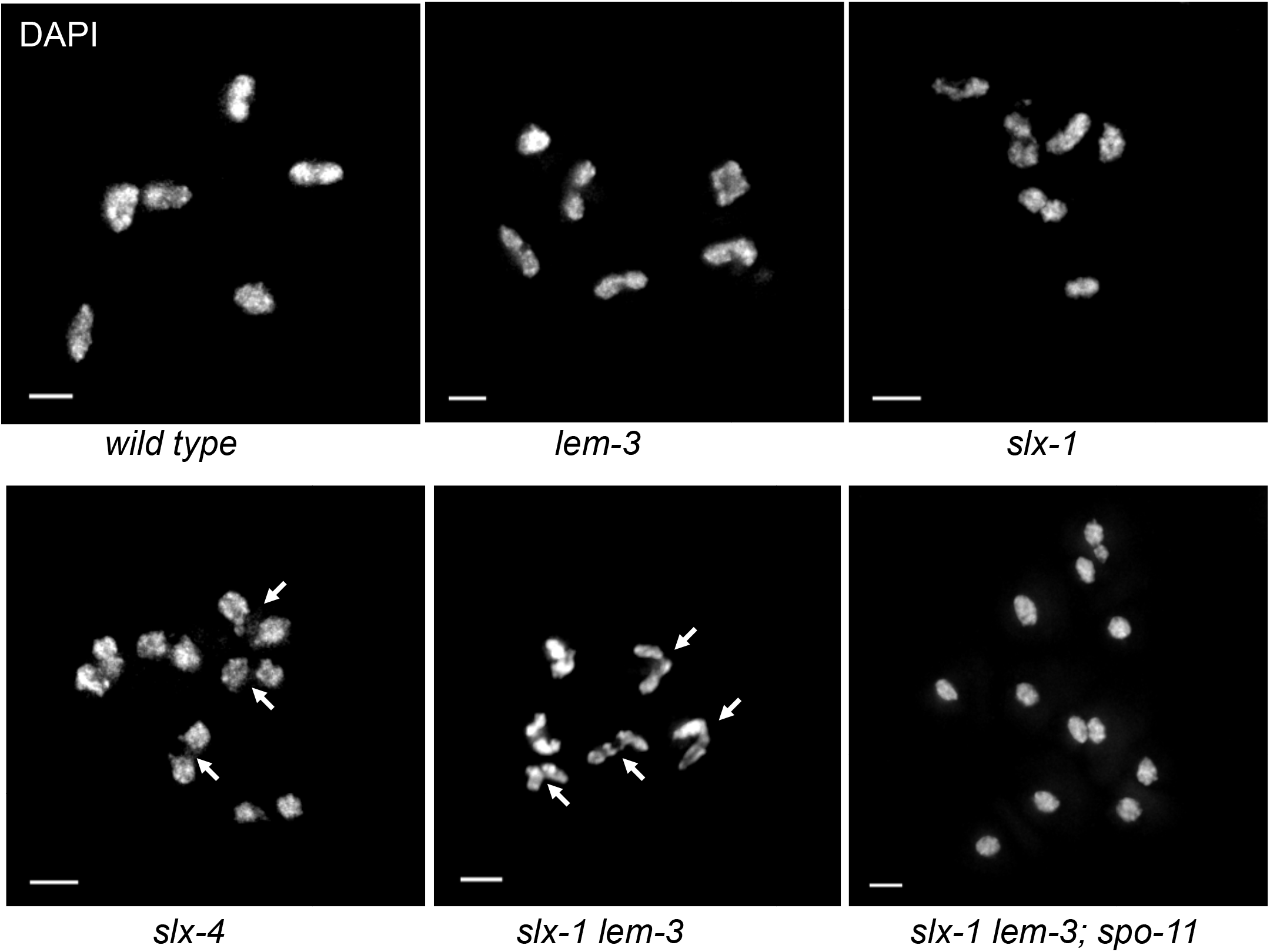
DNA linkages depend on *spo-11* induced meiotic double stranded breaks. White arrows indicate chromosome linkages.

**Figure S3.**
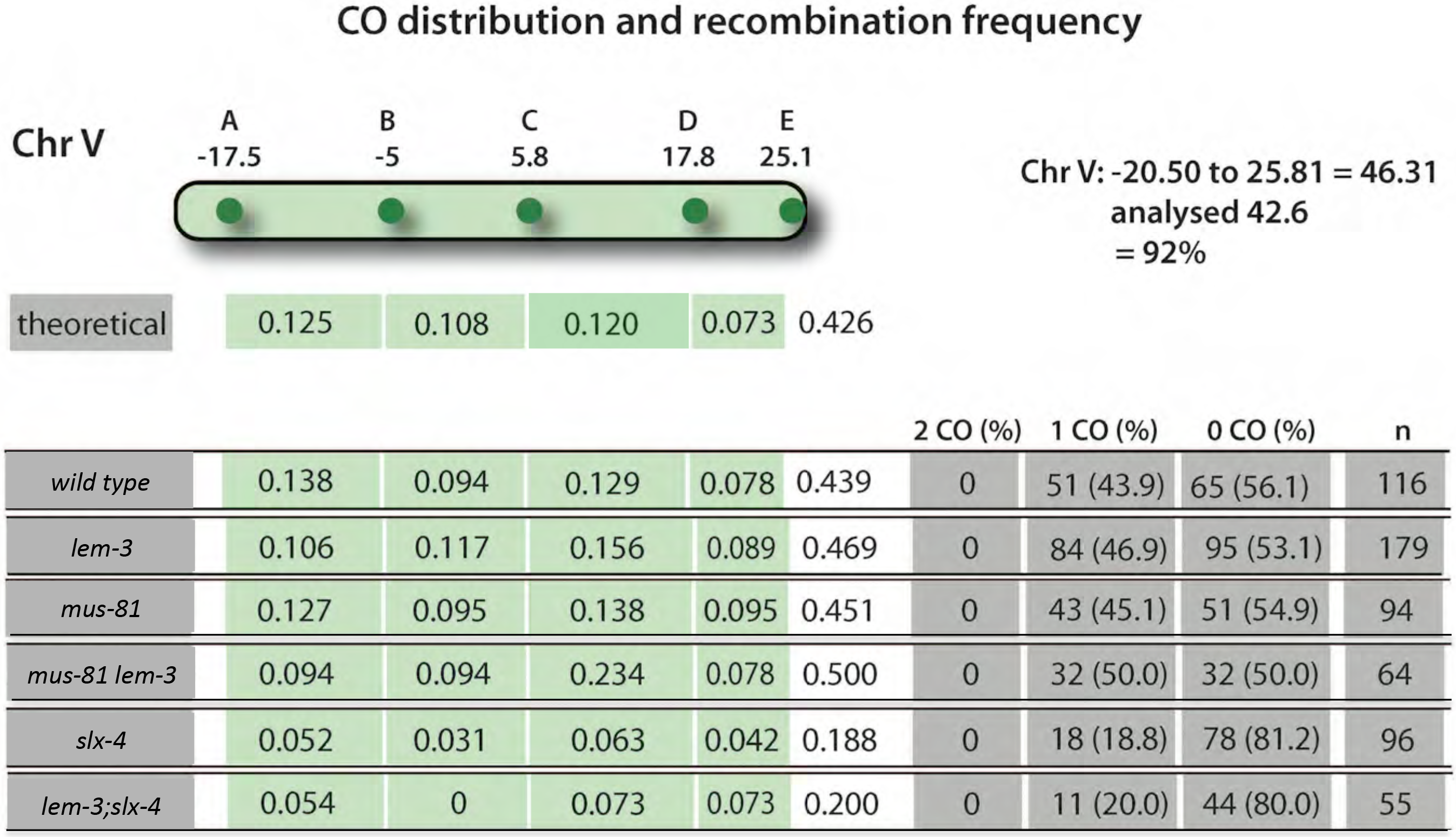
LEM-3 is dispensable for meiotic crossover formation. Analysis of crossover frequencies and distribution on chromosome V. The genetic map positions of the five SNPs, which together cover 92% of chromosome V, are indicated. n is the number of cross-progeny scored. The frequency of 2 COs, 1 CO or 0 CO per chromosome is indicated in absolute numbers and as percentage (in brackets).

**Figure S4.**
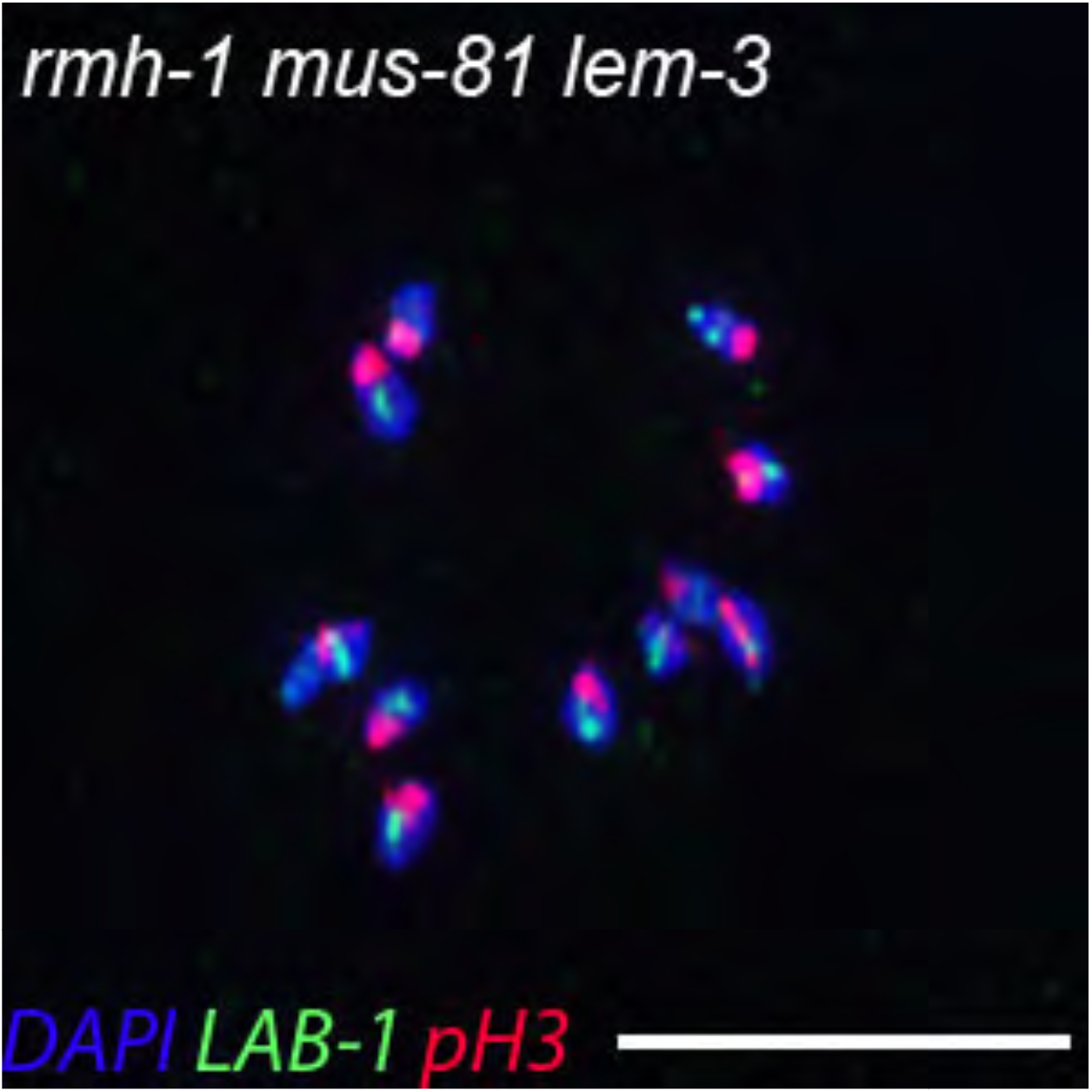
Representative images of peculiar univalent stained by LAB-1 (green) and phsopho histone H3 (pH3, red). Scale bars: 2 μm.

**Figure S5.**
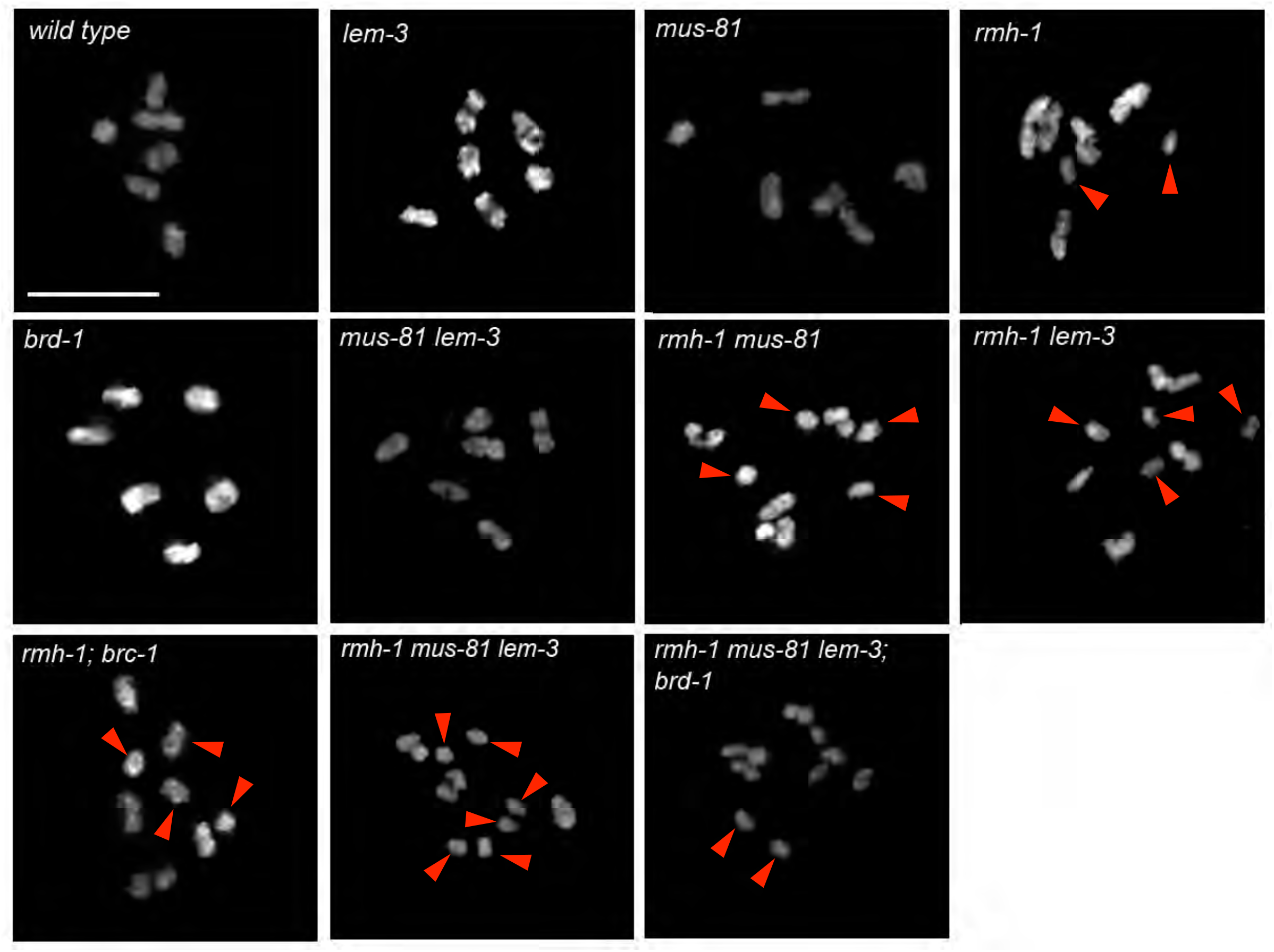
Representative images of DAPI-stained diakinesis chromosomes of indicated genotypes. Univalents are indicated by red arrowheads. Scale bars: 2 μm.

**Movie S1**. Wild type mCherry::H2B, meiosis I and II

**Movie S2**. *lem-3;* mCherry::H2B, meiosis I and II

**Movie S3**. *slx-4;* mCherry::H2B, meiosis I and II

**Movie S4**. *lem-3; slx-4* mCherry::H2B, meiosis I and II

